# Mouse REC114 is essential for meiotic DNA double-strand break formation and forms a complex with MEI4

**DOI:** 10.1101/372052

**Authors:** Rajeev Kumar, Cecilia Oliver, Christine Brun, Ariadna B. Juarez-Martinez, Yara Tarabay, Jan Kadlec, Bernard de Massy

## Abstract

Programmed formation of DNA double strand breaks (DSBs) initiates the meiotic homologous recombination pathway. This pathway allows homologous chromosomes to find each other and the formation of crossing overs, the products of reciprocal exchanges, which are required for proper chromosome segregation at the first meiotic division. Meiotic DSBs are catalyzed by Spo11 that forms a complex with a second subunit, TopoVIBL, and mediates a DNA type II topoisomerase-like cleavage. Several other proteins are essential for meiotic DSB formation, including three evolutionarily conserved proteins first identified in *Saccharomyces cerevisiae* (Mer2, Mei4 and Rec114). These three *S. cerevisiae* proteins and their mouse orthologs (IHO1, MEI4 and REC114) co-localize on the axes of meiotic chromosomes, and mouse IHO1 and MEI4 are essential for meiotic DSB formation. Here, we show that mouse *Rec114* is required for meiotic DSB formation. Moreover, MEI4 forms a complex with REC114 and IHO1 in mouse spermatocytes, consistent with cytological observations. We then demonstrated *in vitro* the formation of a stable complex between REC114 C-terminal domain and MEI4 N-terminal domain. We further determine the structure of REC114 N-terminal domain that revealed similarity with Pleckstrin Homology domains and its property to dimerize. These analyses provide direct insights into the architecture of these essential components of the meiotic DSB machinery.

## Introduction

The conversion from diploid to haploid cells during meiosis requires the expression of a specific and highly differentiated meiotic program in all sexually reproducing eukaryotes. Indeed, meiosis is a specialized cell cycle composed of one replication phase followed directly by two divisions. At the first meiotic division, homologous chromosomes (homologues) are separated through a process called reductional segregation. In most species, reductional segregation requires the establishment of connections between homologues. To achieve this, homologous recombination is induced during meiotic prophase to allow homologues to find each other and to be connected by reciprocal products of recombination (i.e., crossing overs)(Hunter 2015). This homologous recombination pathway is initiated by the formation of DNA double-strand breaks (DSBs) that are preferentially repaired using the homologous chromatid as template (de Massy 2013). Meiotic DSB formation and repair are tightly regulation because DSBs represent a potential threat to genome integrity if they are improperly or not repaired (Sasaki et al. 2010; Keeney et al. 2014). In *Saccharomyces cerevisiae*, several genes are essential for their formation and at least five of them are evolutionarily conserved. *Spo11*, *Top6bl*, *Iho1*, *Mei4* and *Rec114* are the mouse homologs of these five genes *(Baudat et al. 2000; Romanienko and Camerini-Otero 2000; Kumar et al. 2010; Robert et al. 2016; Stanzione et al. 2016)* and are specifically expressed in mouse meiotic cells. SPO11 is orthologous to TopoVIA, the catalytic subunit of archea TopoVI, and is covalently bound to the 5’ ends of meiotic DNA breaks. This indicates that meiotic DSBs are formed by a mechanism with similarity to a type II DNA topoisomerase cleavage (Bergerat et al. 1997; Keeney et al. 1997; Neale et al. 2005). SPO11 acts with a second subunit, TOPOVIBL, orthologous to archaea TopoVIB (Robert et al. 2016; Vrielynck et al. 2016). TOPOVIBL is quite divergent among eukaryotes and in some species, such as *S. cerevisiae*, the orthologous protein (Rec102) shares only one domain of similarity with TOPOVIBL (Robert et al. 2016).

The IHO1, MEI4 and REC114 families have been studied in several organisms, including *S. cerevisiae* (Mer2, Mei4 and Rec114), *Schizosaccharomyces pombe* (Rec15, Rec24 and Rec7), *Arabidopsis thaliana* (PRD3, PRD2 and PHS1), *Sordaria macrospora* (Asy2, Mei4 ortholog not identified, Asy3), *Caenorhabditis elegans* (Mer2 and Mei4 orthologues not identified, DSB-1/2) and *Mus musculus*. Several important properties of these proteins suggest that they act as a complex. Indeed, they colocalize as discrete foci on meiotic chromosome axes in *S. cerevisiae* (Li et al. 2006; Maleki et al. 2007) and *M. musculus* (Stanzione et al. 2016). Their localization is SPO11-independent, as shown for the three *S. cerevisiae* and for the IHO1 and MEI4 *M. musculus* proteins, for *S. pombe* Rec7 (Lorenz et al. 2006) and for *C. elegans* DSB1/2 (Rosu et al. 2013; Stamper et al. 2013). They appear before or at the beginning of meiotic prophase and the number of foci decreases as chromosomes synapse in *S. cerevisiae* (Li et al. 2006; Maleki et al. 2007) and in *M. musculus* (Kumar et al. 2010; Stanzione et al. 2016). In *S. macrospora*, where only Mer2 has been analyzed, its axis localization is also reduced at pachytene upon synapsis (Tesse et al. 2017). In *C. elegans*, foci of the Rec114 orthologs decrease with pachytene progression (Rosu et al. 2013; Stamper et al. 2013). In mice, these foci are on chromosome axes, but they do not colocalize with the DSB repair protein DMC1, also present on axes. This suggests that these foci are displaced upon DSB formation and repair (Kumar et al. 2010). Similarly, chromatin immunoprecipitation (ChIP) experiments in *S. cerevisiae* showed that these three proteins colocalize on chromosome axes, but do not overlap with DSB sites, supporting the hypothesis of a loop/axis interaction for DSB formation (Blat et al. 2002; Panizza et al. 2011). The determinants of their localization are not known, although in *S. cerevisiae* they are detected particularly at domains enriched in Hop1 and Red1, two meiotic-specific axis proteins (Panizza et al. 2011). In addition, several studies have reported the interactions between these three proteins, suggesting a tripartite complex in *S. pombe* (Steiner et al. 2010; Miyoshi et al. 2012), *S. cerevisiae* (Li et al. 2006; Maleki et al. 2007) and *M. musculus* (Kumar et al. 2010; Stanzione et al. 2016). The current knowledge on their *in vivo* direct interactions is limited and based only on yeast two-hybrid assays. Mer2 plays a central role and seems to be the protein that allows the recruitment of Mei4 and Rec114 on chromosome axes. This view is based on the observation that the Mer2 orthologs Rec15 and IHO1, interact with the axis proteins Rec10 (Lorenz et al. 2006) and HORMAD1 (Stanzione et al. 2016) respectively. In *S. cerevisiae*, Mer2 is necessary for Rec114 and Mei4 recruitment to the axis (Panizza et al. 2011). Mer2 is loaded on chromatin before prophase, during S phase, where it is phosphorylated, a step required for its interaction with Rec114 (Henderson et al. 2006; Murakami and Keeney 2014) and for Rec114 recruitment to chromatin (Panizza et al. 2011). Thus, Mer2 coordinates DNA replication and DSB formation. Analysis of the Mer2 ortholog in *S. macrospora* revealed other functions in chromosome structure (Tesse et al. 2017). Overall, it is thought that this putative complex (Mer2/Rec114/Mei4) might directly interact with factors involved in the catalytic activity (i.e., at least Spo11/Rec102/Rec104 in *S. cerevisiae*) at DSB sites. Interactions between Rec114 and Rec102 and Rec104 have been detected by yeast two-hybrid assays (Arora et al. 2004; Maleki et al. 2007). Moreover, in *S. pombe*, an additional protein, Mde2 might bridge the Mer2/Rec114/Mei4 and Rec12/Rec6/Rec14 complexes (Miyoshi et al. 2012). The direct implication of the Mer2/Mei4/Rec114 complex in DSB activity is also supported by its detection at DSB sites (Panizza et al. 2011; Miyoshi et al. 2012). However, no specific feature or domain has been identified in Mei4 or Rec114 to understand how they may regulate DSB activity. One could hypothesize that they play a direct role in activating or recruiting the Spo11/TopoVIBL complex for DSB formation. The hypothesis that these proteins might regulate DSB formation through some interactions is also consistent with the findings that Rec114 overexpression inhibits DSB formation in *S. cerevisiae* (Bishop et al. 1999) and that altering Rec114 phosphorylation pattern can up- or down- regulate DSB levels (Carballo et al. 2013). It is possible that Rec114 and Mei4 have distinct roles, because Spo11 non-covalent interaction with DSBs is Rec114-dependent but Mei4-independent (Prieler et al. 2005), and Spo11 self-interaction depends on Rec114 but not on Mei4 (Sasanuma et al. 2007). However, in *Zea mays* and *A. thaliana*, the *Rec114* ortholog (*Phs1*) seems not to be required for DSB formation (Pawlowski et al. 2004; Ronceret et al. 2009). Here, we performed a functional and molecular analysis to determine whether mouse REC114 is required for meiotic DSB formation, and whether it interacts directly with some of its candidate partners.

## Results

### Rec114-null mutant mice are deficient in meiotic DSB formation

We analyzed mice carrying a null allele of *Rec114*. In the mutated allele (here named Rec114^-^ and registered as Rec114^tm1(KOMP)Wtsi^) exon 3 and 4 were deleted and a lacZ-neomycin cassette was inserted upstream of this deletion (Fig. 1A; Supplemental Fig. S1A, S1B). This allele may encode a truncated protein of 18KD without the conserved motifs SSM3, 4, 5 and 6 (Supplemental Fig. S1B) (Kumar et al. 2010; Tesse et al. 2017). We also generated from this allele, another null allele (named and registered as *Rec114^del^*) without the insertion cassette. We performed all subsequent analyses using mice with the *Rec114^-^* allele unless otherwise stated, and confirmed several phenotypes in mice carrying the *Rec114^del^* allele. Heterozygous (*Rec114^+/-^*) and homozygous (*Rec114^-/-^*) mutant mice were viable. We confirmed the absence of REC114 protein in *Rec114^-/-^* mice by western blot analysis of total testis extracts and after REC114 immunoprecipitation and could not detected the putative truncated protein expressed from the *Rec114*^-^ allele around 18KD (Fig. 1B and data not shown).

**Figure 1.**
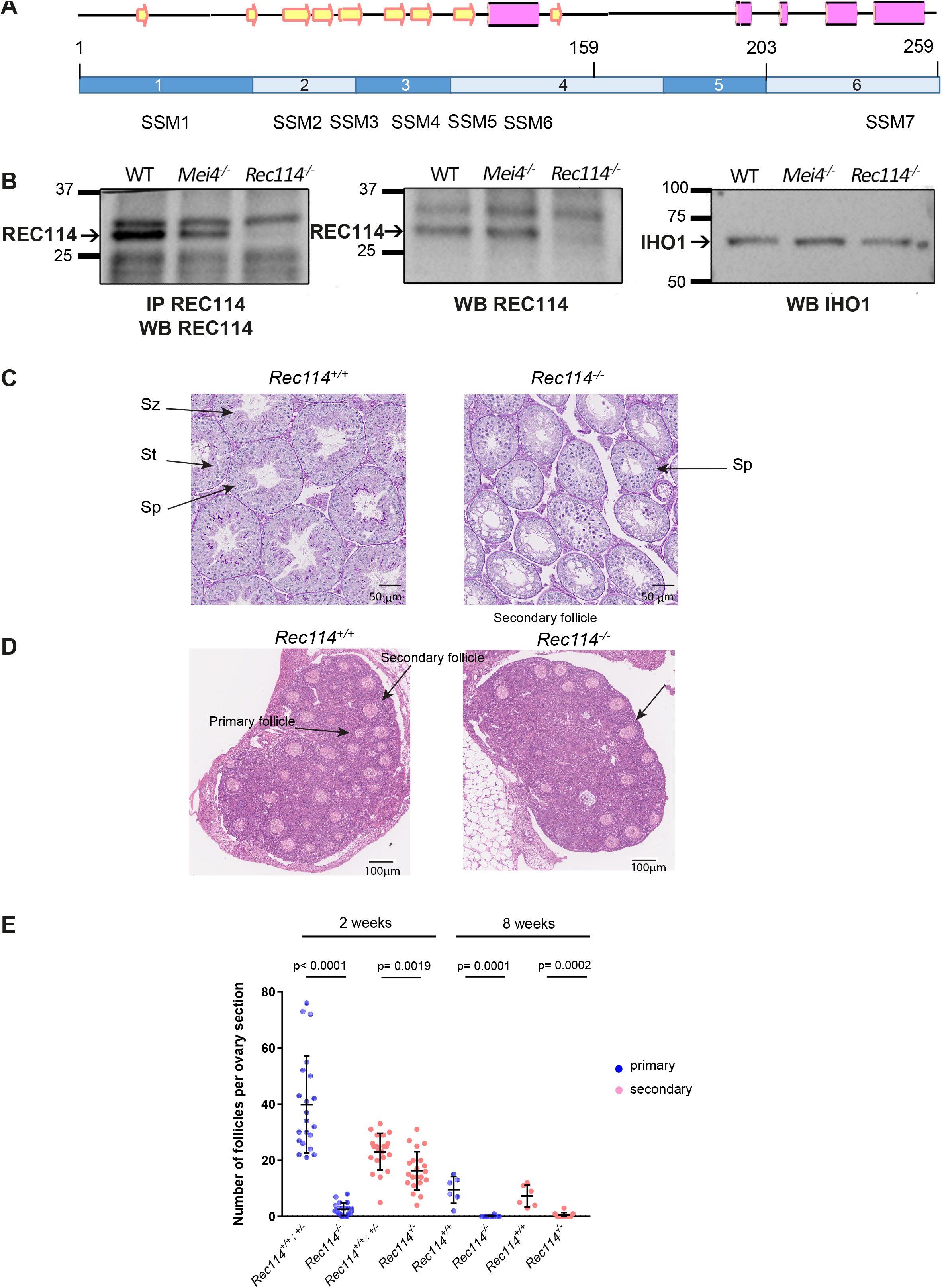
REC114 is essential for spermatogenesis and oogenesis. (A) Conserved domains and organization of REC114. The conserved motifs are SSM1 to 7 (Kumar et al. 2010). Secondary structures were predicted with PSIPRED v3.3 (http://bioinf.cs.ucl.ac.uk/psipred/). Pink cylinders, α–helices; yellow arrows, β–sheets. Exons (1 to 6) are shown as dark and light blue rectangles. (B) REC114 is not detected in *Rec114^-/-^* mice. Western blot (WB) analysis of total testis extracts from wild type (WT), *Mei4^-/-^* and *Rec114^-/-^* as control, prepubertal mice (14 days post-partum, dpp) with anti-REC114 (central panel), with anti-IHO1 (right panel) and anti-REC114 antibodies after immunoprecipitation of REC114 (left panel). (C) Spermatogenesis is defective in *Rec114^-/-^* mice Periodic Acid Schiff staining of testis sections from 9-week-old *Rec114^+/+^* and *Rec114^-/-^* mice. Sz: spermatozoa; St: round spermatid; Sp: spermatocyte. (D) Oogenesis is defective in *Rec114^-/-^* mice Hematoxylin-eosin staining of ovary sections from 2-week-old *Rec114^+/+^* and *Rec114^-/-^* mice. (E) Quantification of primary and secondary follicles in ovaries from 2-week-old and 8-week-old *Rec114^+/+^*, *Rec114^+/-^* and *Rec114^-/-^* mice. At 2 weeks of age, the numbers (mean ± SD) of primary (blue circles) and secondary (pink circles) follicles were 39.9±17.3 and 23.1±6.6, respectively, for *Rec114^+/+^* (n=3) and *Rec114^+/-^* (n=1) mice (*Rec114^+/+^* and *Rec114^+/-^* data were pooled, n sections=21 in total) and 2.6±2.2 and 16.3±6.9, respectively, for *Rec114^-/-^* mice (n=5; n sections=21). At 8 weeks of age, the numbers (mean ± SD) of primary (blue circles) and secondary (pink circles) follicles were 9.5±4.8 and 7.3±3.8, respectively, for *Rec114^+/+^* mice (n=1; n sections=6), and 0.1±0.3 and 0.5±1.0, respectively, for *Rec114^-/-^* mice (n=2; n sections= 10). P values were calculated with the Mann-Whitney two-tailed test.

To monitor the consequences of REC114 absence on gametogenesis, we performed histological analysis of testes and ovaries. Spermatogenesis was altered in *Rec114^-/-^* adult male mice, as indicated by the presence of major defects in testis tubule development compared with wild type (*Rec114^+/+^*) mice (Fig. 1C). Specifically, in *Rec114^-/-^* animals the tubule diameter was smaller and tubules lacked haploid cells (spermatids and spermatozoa). In these tubules, the most advanced cells were spermatocytes, although some were also depleted of spermatocytes. Testis weight was significantly lower in *Rec114^-/-^* than wild type mice (Supplemental Fig. S1C). In ovaries from *Rec114^-/-^* mice, oogenesis was significantly affected, as indicated by the strongly reduced number of primary and secondary follicles at two weeks post-partum and their nearly absence at eight weeks (Fig. 1D, E). Consistent with these gametogenesis defects, *Rec114^-/-^* males and females were sterile. Indeed, mating of wild type C57BL/6 animals with *Rec114^-/-^* males and females (n=3/sex) crossed for four months yielded no progeny.

To investigate the nature of the meiotic defect, we monitored by cytological analysis the presence and localization of various markers of recombination and homologous chromosome interactions during meiotic prophase. The formation of meiotic DSBs was followed by the detection of γH2AX, the phosphorylated form of H2AX which is enriched in chromatin domains around DSB sites. The DSB repair activity was assessed by the detection of the strand exchange proteins RAD51 and DMC1 and of a subunit (RPA2) of the single strand DNA binding protein complex RPA. Chromosome axes and assembly of the synaptonemal complex were monitored by detection of SYCP3 and SYCP1 respectively. Detection of γH2AX revealed that in *Rec114^-/-^* mice, meiotic DSBs were absent or strongly reduced in both spermatocytes and oocytes, whereas chromosome axes formed normally, based on SYCP3 detection (Fig. 2A). Quantification of the γH2AX signal indicated a 16- and 11-fold reduction in *Rec114^-/-^* spermatocytes and oocytes, respectively, compared with wild type gametocytes (Fig. 2B). Consistent with this defect in DSB formation, DSB repair foci were strongly reduced. Specifically, foci of DMC1 were reduced in *Rec114^-/-^* spermatocytes and oocytes compared with wild type cells (Fig. 2C, 2D; Supplemental Fig. S2A, S2B). Similarly, RPA2 and RAD51 foci were strongly reduced or undetectable in *Rec114^-/-^* compared with wild type gametocytes (Supplemental Fig. S2C, S2D). Meiotic DSB formation and repair promotes interactions between homologues that are stabilized by the loading of SYCP1, a component of the synaptonemal complex (Fraune et al. 2012). Analysis of SYCP1 localization during meiosis showed major defects in both male and female *Rec114^-/-^* meiocytes. The presence of short SYCP1 stretches suggested progression into zygonema; however, these stretches never elongated to form a full length synaptonemal complex between homologues, indicating failure of homologous synapsis formation (Fig. 2E and Supplemental Fig. S2E). Altogether, the phenotypes of *Rec114^-/-^* mice are highly similar to those of the previously characterized *Spo11^-/-^* (Baudat et al. 2000; Romanienko and Camerini-Otero 2000), *Mei1^-/-^* (Libby et al. 2002; Libby et al. 2003), *Mei4^-/-^* (Kumar et al. 2010), *Iho1^-/-^* (Stanzione et al. 2016) and *Top6bl^-/-^* (Robert et al. 2016) mice where the formation of DSBs, of DSB repair foci and of homologous synapses is strongly affected. Histological and cytological analyses of *Rec114^del/del^* mutant mice showed similar phenotypes, indicating that the cassette present in the *Rec114^-^* allele does not cause the observed meiotic defects (Supplemental Fig. S3A-E).

**Figure 2.**
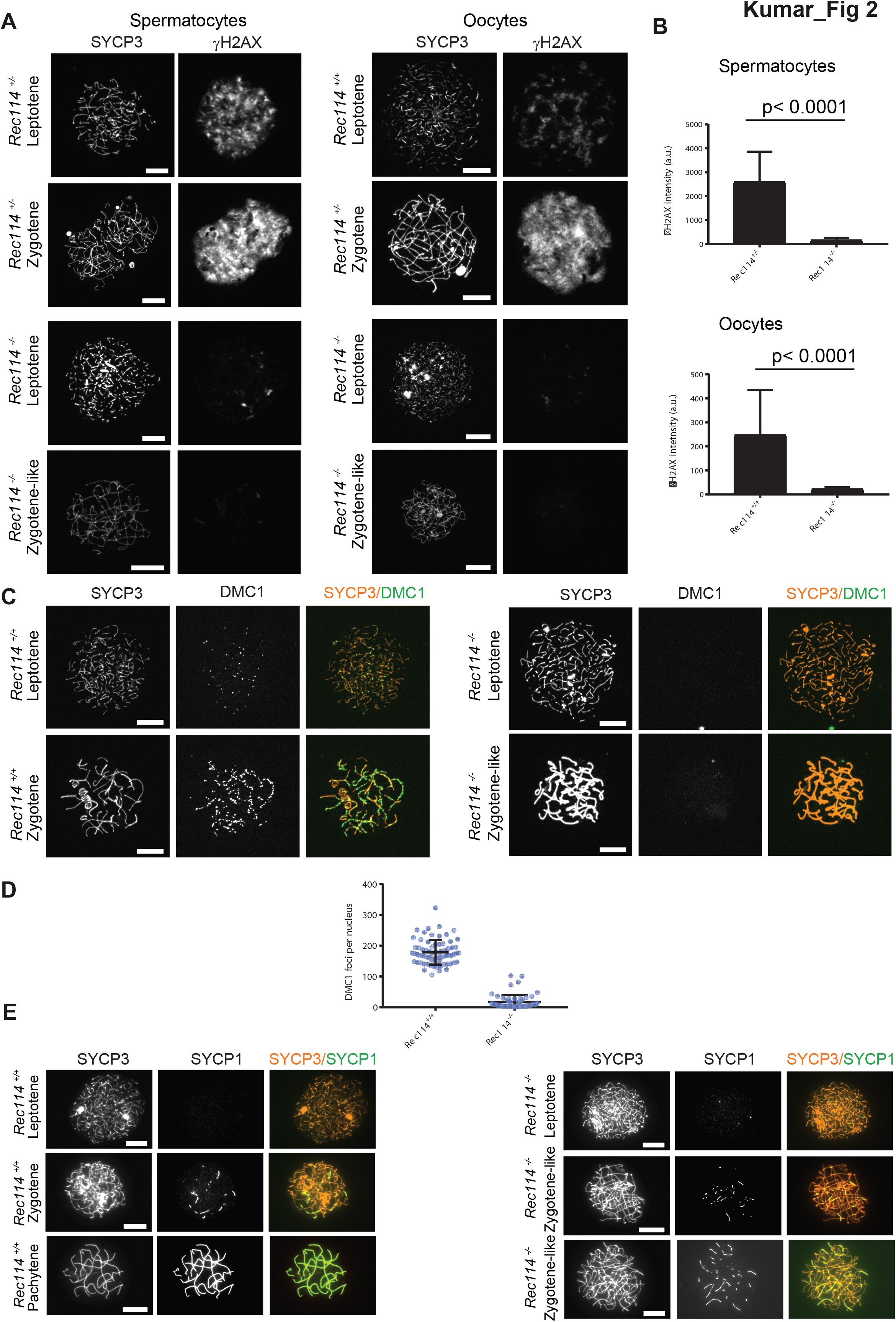
*Rec114^-/-^* mice show defects in DSB formation and homologous synapsis. (A) Immunostaining of γH2AX and SYCP3 in spermatocytes from 13 dpp *Rec114^+/-^* and *Rec114^-/-^* males, and from E15 (15 days of embryonic development) *Rec114^+/+^*, *Rec114^+/-^* and *Rec114^-/-^* oocytes. In *Rec114^-/-^* spermatocytes and oocytes, no pachynema could be observed and spermatocytes or oocytes with partially synapsed chromosomes were defined as zygotene-like. Scale bar, 10μm. (B) Quantification of the total γH2AX signal per nucleus (mean ± SD; a.u: arbitrary units) on spreads from leptotene spermatocytes (13 dpp) and from leptotene oocytes (E15): 2597±1261 and 165±95 in *Rec114^+/-^* and *Rec114^-/-^* males, respectively (n= 53 and 50); 248±187 and 23±7 in *Rec114^+/+^* and *Rec114^-/-^* females, respectively (n= 48 and 47).. P values were calculated with the Mann-Whitney two-tailed test. (C) Immunostaining of DMC1 and SYCP3 in spermatocytes from 15dpp *Rec114^+/+^* and *Rec114^-/-^* males. Scale bar, 10μm. (D) Quantification of DMC1 foci (mean ± SD) in leptotene and zygotene spermatocytes from *Rec114^+/+^* and *Rec114^-/-^* mice (178.6±39.9 and 17.2±23.0 in *Rec114^+/+^* and *Rec114^-/-^* males, respectively; n= 71 and 59). P < 0.0001 (Mann-Whitney two-tailed test). (E) Immunostaining of SYCP1 and SYCP3 in oocytes from *Rec114^+/+^* females (E15, E15 and E16 for leptotene, zygotene and pachytene, respectively) and *Rec114^-/-^* females (E15, E16 and E17 for leptotene, zygotene-like and zygotene-like, respectively). Scale bar, 10μm.

### In vivo REC114 interacts with MEI4 and these proteins display a mutually dependent localization

REC114, MEI4 and IHO1 colocalize on the axis of meiotic chromosomes, and IHO1 is needed for MEI4 loading (Stanzione et al. 2016). First, we tested whether IHO1 loading required REC114 and MEI4. This was clearly not the case because IHO1 localization was similar in wild type and in *Rec114^-/-^* and *Mei4^-/-^* spermatocytes (Fig. 3A and Supplemental Fig. S4A). This observation is consistent with a role for IHO1 in REC114 and MEI4 recruitment.

**Figure 3.**
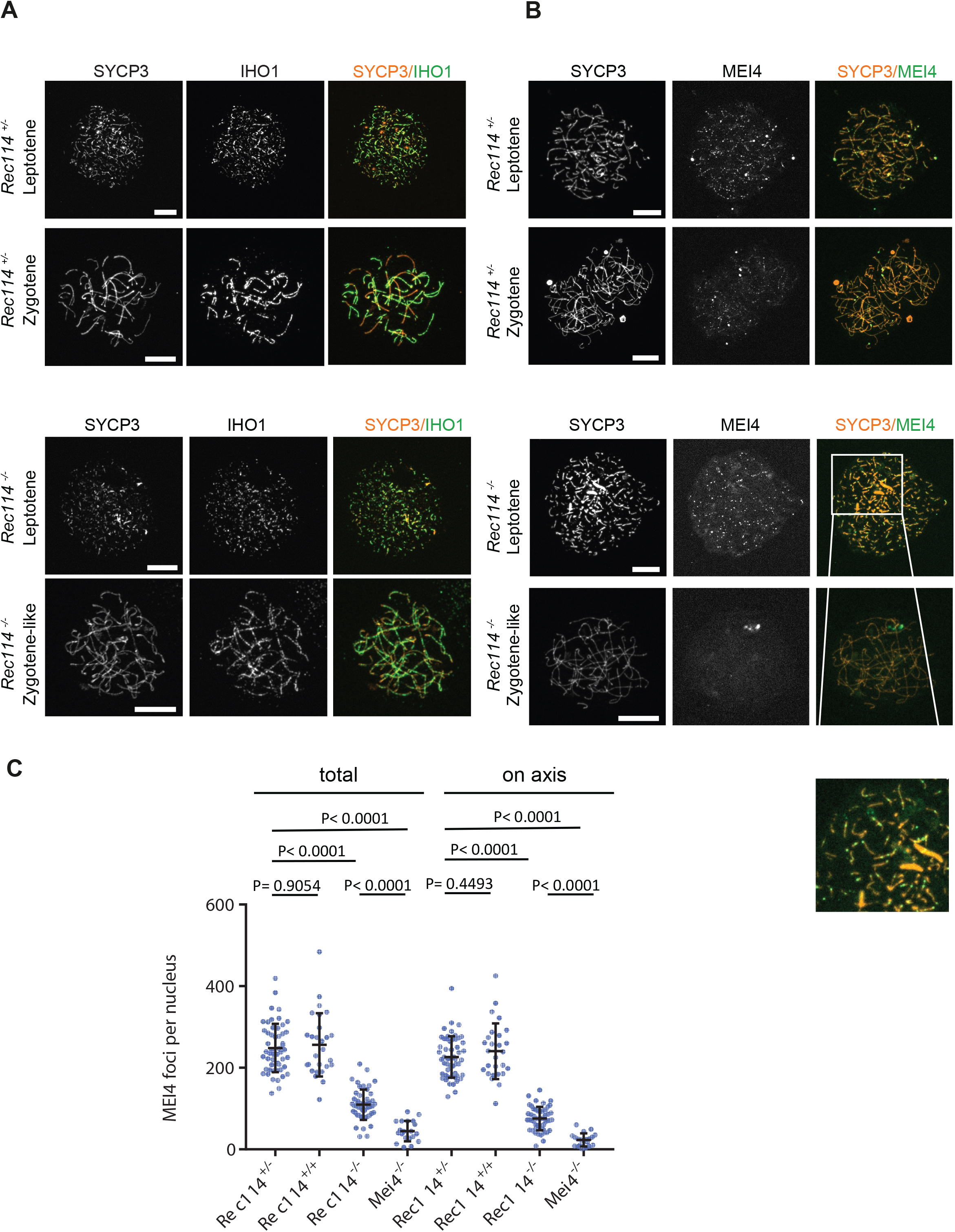
REC114 is required for robust MEI4 foci localization. (A) Immunostaining of IHO1 and SYCP3 in early prophase spermatocytes from 13 dpp *Rec114^+/-^* and *Rec114^-/-^* males. Scale bar, 10μm. (B) Immunostaining of MEI4 and SYCP3 in early prophase spermatocytes from 13 dpp *Rec114^+/-^* and *Rec114^-/-^* males. Scale bar, 10μm. (C) Quantification of MEI4 foci in leptotene spermatocytes from 13 dpp *Rec114^+/-^*, *Rec114^+/+^*, *Rec114^-/-^* and *Mei4^-/-^* males. The numbers (mean ± SD) of total foci and of foci on chromosome axes (colocalized with SYCP3) were: 248±59 and 226±51 for *Rec114^+/-^* (n=53 nuclei), 256±78 and 241±68 for *Rec114^+/+^* (n=27), 109±37 and 76±29 for *Rec114^-/-^* (n=50), 45±25 and 23±16 for *Mei4^-/-^* (n=20), respectively. P values were calculated with the Mann-Whitney two-tailed test.

We then tested whether MEI4 and REC114 regulated each other localization. MEI4 forms 200-300 foci on meiotic chromosome axes at leptonema. Then, the focus number progressively decreases as cells progress into zygonema and MEI4 becomes undetectable at pachynema. This expression decrease during meiotic progression is directly correlated with synapsis formation (MEI4 foci are specifically depleted from synapsed axes) and with DSB repair (MEI4 foci are excluded from DMC1 foci) (Kumar et al. 2010). At leptotene, the number of axis-associated MEI4 foci was reduced by 3- to 4-fold in *Rec114^-/-^* spermatocytes and oocytes (Fig. 3B, 3C; Supplemental Fig. S4B, S4C) and their intensity was significantly decreased (by 1.75-fold in spermatocytes and by 1.9-fold in oocytes) compared with wild type controls (Supplemental Fig. S4D). The MEI4 signal detected in *Rec114^-/-^* gametocytes was higher than the non-specific background signal observed in *Mei4^-/-^* spermatocytes (Fig. 3C). This suggests that REC114 contributes, but it is not essential for MEI4 focus formation on meiotic chromosome axis.

REC114 foci colocalize with MEI4 and, like MEI4 foci, their number is highest at leptonema and then progressively decreases upon synapsis (Stanzione et al. 2016). We thus tested whether REC114 foci required MEI4 for axis localization. At leptotene, few axis-associated REC114 foci above the background signal could be detected in *Mei4^-/-^* spermatocytes, where their number was reduced by more than 10-fold compared with wild type cells (Fig. 4A, B). However, this low level of REC114 foci in *Mei4^-/-^* was still significantly higher compared with the number in *Rec114^-/-^* gametocytes (Fig. 4B). REC114 foci were not reduced in *Spo11^-/-^* mice (Fig. 4B), as previously reported for MEI4 foci (Kumar et al. 2010). This indicates that these proteins are loaded on the chromosome axis independently of SPO11 activity. Overall, MEI4 and REC114 are reciprocally required for their localization.

**Figure 4.**
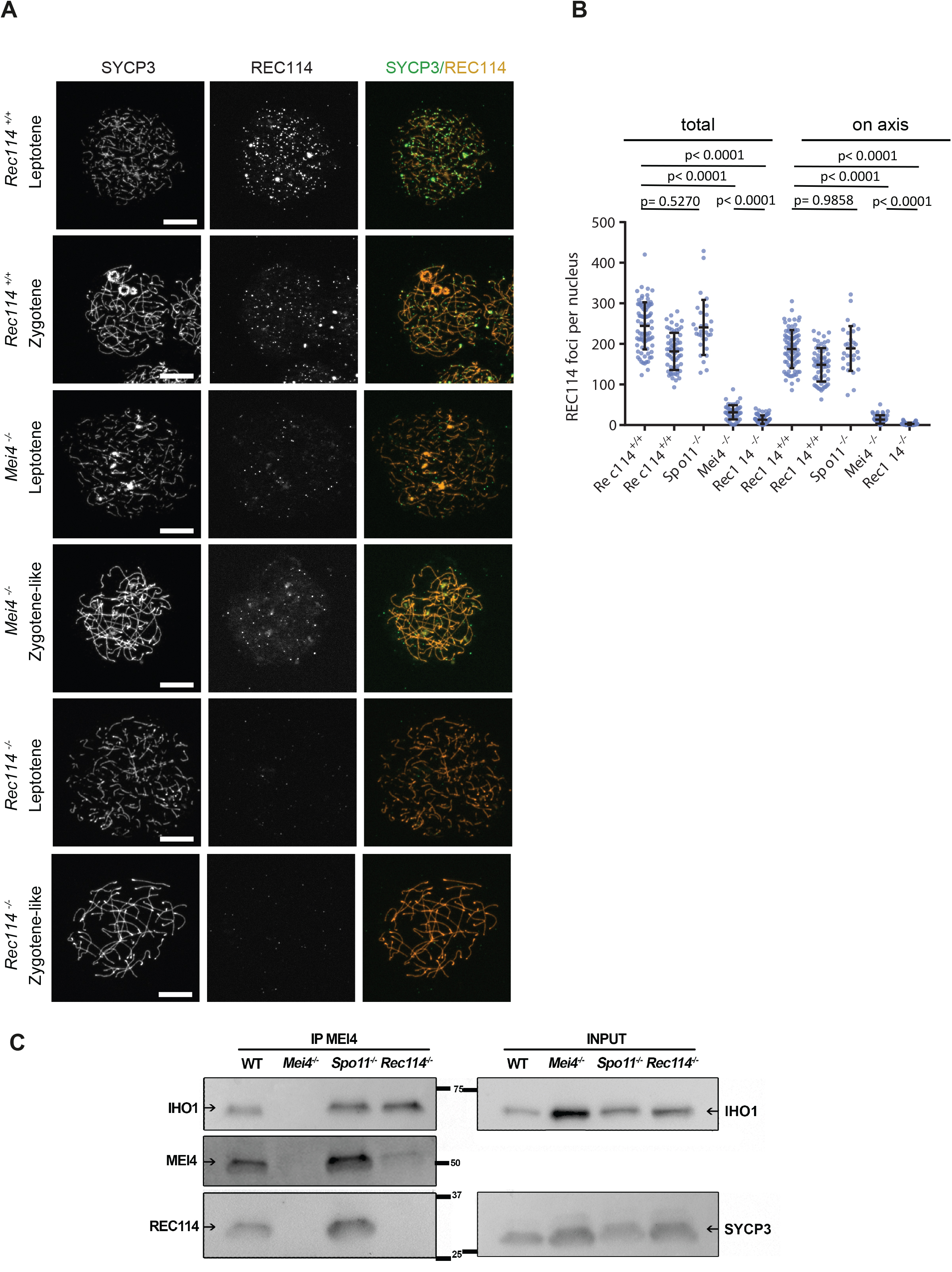
MEI4, REC114 and IHO1 form a complex. (A) Immunostaining of REC114 and SYCP3 in early prophase spermatocytes from *Rec114^+/+^* (14dpp), *Mei4^-/-^* (12dpp) and *Rec114^-/-^* (14dpp) males. Scale bar, 10μm. (B) Quantification of REC114 foci in leptotene spermatocytes from 13 dpp *Rec114^+/+^*, *Mei4^-/-^*, *Spo11^-/-^* and *Rec114^-/-^* males. Two independent *Rec114^+/+^* spread preparations were use as controls for *Mei4^-/-^* and *Rec114^-/-^* samples. The numbers (mean ± SD) of total foci and of foci on chromosome axis (colocalized with SYCP3) were: 245±58 and 187±47 for *Rec114^+/+^* (n=83 nuclei), 181±46 and 149±41 for *Rec114^+/+^* (n=63), 241±68 and 189±55 for *Spo11^-/-^* (n=30), 13±11 and 3±3 for *Rec114^-/-^* (n=69), 32±17 and 14±11 for *Mei4^-/-^* (n=52). P values were calculated with the Mann-Whitney two-tailed test. (C) Co-immunoprecipitation of REC114 and IHO1 with MEI4. Total testis extracts from 14 ddp *Rec114^+/+^* (WT), *Mei4^-/-^*, *Spo11^-/-^* and *Rec114^-/-^* mice were immune-precipitated with an anti-MEI4 antibody. Input extracts were probed with anti-IHO1 and anti-SYCP3 antibodies. Immunoprecipitated fractions were probed with anti-IHO1, anti-MEI4 and anti-REC114 antibodies.

MEI4 and REC114 colocalization, their mutual dependency for robust localization and their interaction in yeast two-hybrid assays (Kumar et al. 2010) strongly suggested that these two proteins interact directly or indirectly *in vivo*. Indeed, we could detect REC114 after immunoprecipitation of MEI4 (Fig. 4C). Interestingly, IHO1 also was immunoprecipitated, in agreement with the cytological analysis, suggesting that MEI4 interacts with both IHO1 and REC114. These three proteins could be part of the same complex, or form two independent complexes. Although MEI4 and IHO1 did not interact in a yeast two-hybrid assay (Stanzione et al. 2016), the detection of IHO1 after immunoprecipitation of MEI4 in *Rec114^-/-^* extracts (Fig. 4C) suggests a direct or indirect interaction between IHO1 and MEI4 in mouse spermatocytes. This observation is consistent with the low number, but above background, of MEI4 foci on chromosome axes detected in *Rec114^-/-^* gametocytes (Fig. 3B, 3C). Immunoprecipitation experiments with an anti-REC114 antibody allowed the detection of REC114, but the MEI4 or IHO1 signals were too weak to draw clear conclusions (data not shown).

### REC114 and MEI4 form a stable complex

These *in vivo* assays suggested that REC114 and MEI4 directly interact. To verify whether they form a stable complex, we produced recombinant full length REC114 in bacteria (Fig. 5). However, we could not produce recombinant full length MEI4 or its N-terminal or C-terminal domains alone. Conversely, when co-expressed with REC114, the N-terminal fragment (1-127) of MEI4 was soluble and could be co-purified with REC114 on Strep-Tactin resin (Fig. 5A, lane 4), providing the first evidence of a direct interaction between REC114 and MEI4. To identify the REC114 region that interacts with MEI4, we produced the N-terminal domain and a C-terminal fragment (residues 203-254) of REC114 and found that the REC114 C-terminal region (but not the N-terminal domain) was sufficient for binding to MEI4 (Fig. 5A, lanes 5, 6). Finally, we could purify the MEI4 (1-127) and REC114 (203-254) complex and show that the two proteins co-eluted as a single peak from the Superdex 200 gel filtration column (Fig. 5B, C).

**Figure 5.**
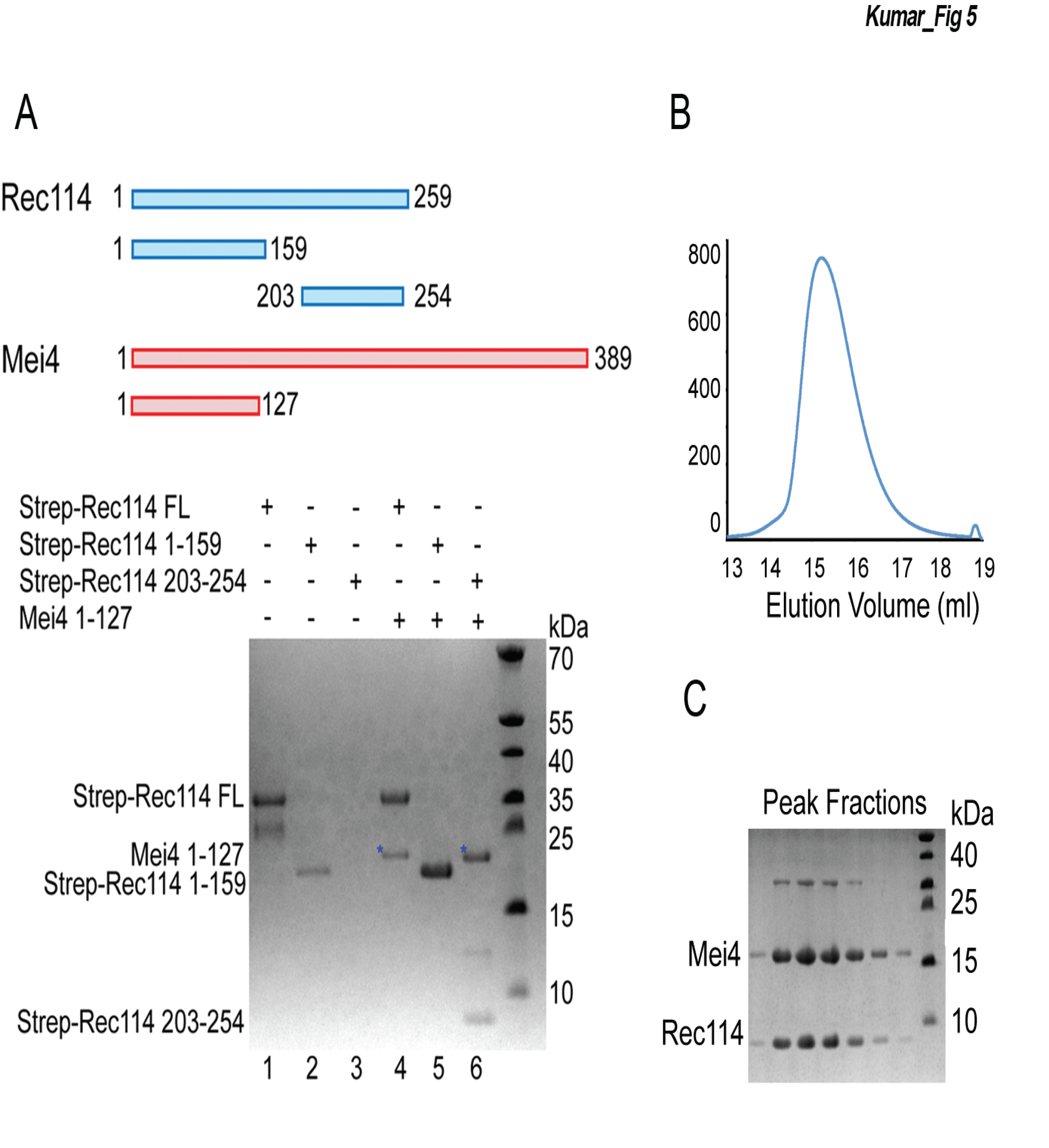
REC114 forms a stable complex with MEI4. (A) Strep-tag pull-down experiments with the REC114 and MEI4 constructs shown in the upper panel. Full length and fragments of Strep-REC1114 were purified alone or after co-expression with MEI4 (1-127). Strep-REC114 (203-254) was insoluble on its own (lane 3), but when co-expressed with MEI4 (1-127) became soluble and was sufficient for interaction with MEI4 (lane 6). MEI4 (1-127, blue star) was pulled down by full length (FL) REC114 and REC114 (203-254), but not by REC114 (1-159)(lanes 4-6). Proteins were detected by Coomassie blue staining. (B) Strep-tagged REC114 (203-254) was co-expressed with MEI4 (1-127) and purified first using Strep-Tactin resin, and then Superdex 200 size-exclusion chromatography. The gel filtration elution profile is shown. (C) SDS-PAGE analysis of the peak fractions shown in B. Proteins were detected by Coomassie blue staining.

### Rec114 contains a Pleckstrin homology domain

To gain insights into the structure of mouse REC114 we produced the full length protein in bacteria. Then, using limited trypsin proteolysis we identified a stable fragment (residues 15-159) that was suitable for structural analysis. We determined the crystal structure of this REC114 N-terminal region at a resolution of 2.5 Å by SAD using the selenomethionine (SeMet)-substituted protein. The final model, refined to an *R*_free_ of 30% and *R*-factor of 25% included residues 15-150 (Table 1).

**Table 1.** Data collection and refinement statistics for REC114 (15-159).

|  | **REC114 native** | **REC114**<br>**SeMet** |
| --- | --- | --- |
| **Data collection** |  |  |
| Space group | *P*6<sub>1</sub>22 | *P*4<sub>2</sub>2<sub>1</sub>2 |
| Cell dimensions |  |  |
| *a*, *b*, *c* (Å) | 107.5 107.5 82.8 | 88.5 88.5 101.5 |
| $\alpha$ $\beta$ $\gamma$ (°) | 90, 90, 120 | 90, 90, 90 |
| Resolution (Å) | 93-2.5 (2.57-2.5)<sup>a</sup> | 66.7-2.7 (2.8-2.7) |
| *R*<sub>merge</sub> | 5.7 (212.1) | 12.2 (108.8) |
| I / σI | 24.2 (1.11) | 11.1 (1.68) |
| *CC*<sub>1/2</sub> | 1 (0.548) | 0.997 (0.725) |
| Completeness (%) | 99.6 (97.5) | 100 (99.9) |
| Redundancy | 11.2 (11.1) | 7.1 (7.1) |
| **Refinement** |  |  |
| Resolution (Å) | 46.6-2.5 |  |
| No. reflections | 10215 |  |
| *R*<sub>work</sub>/*R*<sub>free</sub> | 25(30) |  |
| *B*-factors | 77.3 |  |
| R.m.s. deviations |  |  |
| Bond lengths (Å) | 0.014 |  |
| Bond angles (°) | 1.641 |  |

<sup>a</sup> Values in parentheses are for the highest resolution shell.

Unexpectedly, the structure revealed that REC114 (15-150) forms a pleckstrin homology (PH) domain, with two perpendicular antiparallel β-sheets followed by a C-terminal helix (Fig. 6). Several residues are disordered in loops between the β strands. In the SeMet protein dataset that we solved at 2.7 Å, the crystallographic asymmetric unit contained two REC114 molecules, but the position of β2 that packs against β1 in one of the molecules was shifted by three residues. A Protein Data Bank search using the PDBeFold server at EBI revealed that REC114 (15-150) was highly similar to other PH domains and that the N-terminal domain of the CARM1 arginine methyltransferase (PDB code 2OQB) was the closest homolog (Supplemental Fig. S5).

**Figure 6.**
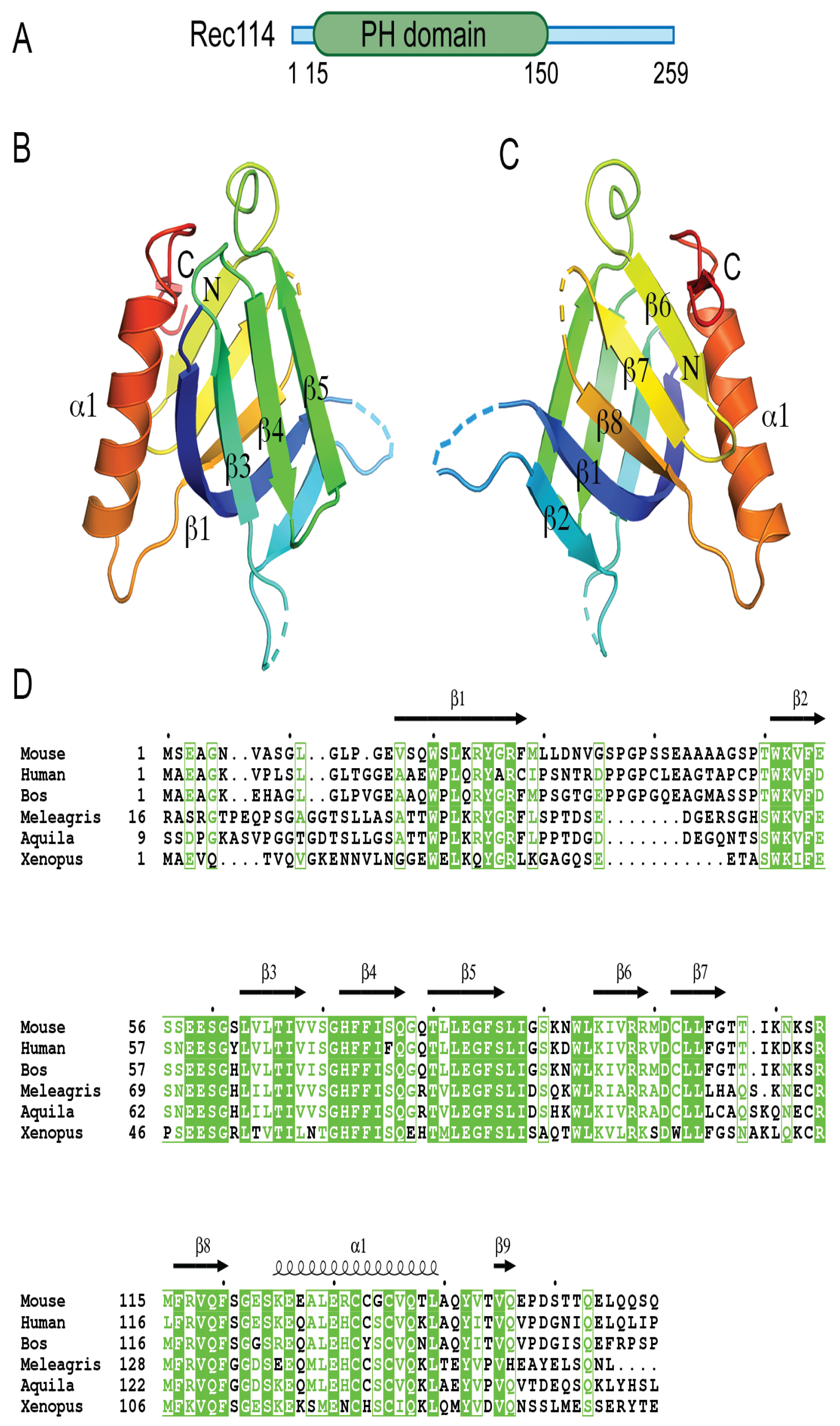
Crystal structure of the REC114 pleckstrin homology (PH) domain. (A) Schematic representation of the PH domain structure in mouse REC114, based on this work. (B) Ribbon diagram of the REC114 PH domain. The polypeptide chain is colored from the N terminus (blue) to the C terminus (red). Missing residues in the loops are shown by dashed lines. (C) The same ribbon diagram as in A, but rotated 180° around the vertical axis. (D) Sequence alignment of REC114 proteins. Residues that are 100% conserved are in solid green boxes. The secondary structures of REC114 are shown above the sequences. Mouse, *Mus musculus*: NP_082874.1; Human, *Homo sapiens*: NP_001035826.1; Bos, *Bos mutus*: XP_005907161.1; Meleagris, *Meleagris gallopavo*: XP_019474806.1; Aquila, *Aquila chrisaetos canadensis*: XP_011595470.1; Xenopus, *Xenopus laevis*: OCT89407.1.

Mapping the conserved residues to the protein surface revealed that both β-sheets contained exposed conserved residues that could be involved in protein interactions with REC114 partners (Supplemental Fig. S6). In the crystal, the PH domain formed extensive crystallographic contacts with a symmetry-related molecule. Indeed, this interface, judged as significant using the PDBePisa server, buried a surface of 764 Å^2^ and included several salt bridges and hydrogen bonds formed by the well-conserved Arg98 of β6 and Glu130 and Gln137 of α1 (Supplemental Fig. S7A). To test whether REC114 dimerized in solution, we analyzed the PH domain by size exclusion chromatography-multiple angle laser light scattering (SEC-MALLS) that allows measuring the molecular weight. Although the monomer molecular mass of the fragment was 16 kDa (32 kDa for a dimer), the MALLS data indicated a molecular weight of 24.7 kDa for the sample at the concentration of 10 mg/ml. When injected at lower concentrations, the protein eluted later and the molecular weight diminished (21 kDa at 5 mg/ml) (Supplemental Fig. S7B). These results could be explained by a concentration-dependent dimerization of the PH domain with a fast exchange rate between monomers and dimers that co-purify together during SEC. The physiological significance of possible REC114 dimers requires additional investigations.

## Discussion

Previous studies in yeast have shown that the putative complex involving *S. cerevisiae* Rec114, Mei4 and Mer2 is essential for meiotic DSB formation. Their localization at DSB sites (observed by ChIP) suggests that in yeast, this complex may play a direct role in promoting DSB activity. Several studies have shown the evolutionary conservation of these three partners. In mammals, MEI4 and IHO1 (the Mei4 and Mer2 orthologs, respectively) are required for meiotic DSB formation (Kumar et al. 2010; Stanzione et al. 2016). Here, we show that REC114 function in the formation of meiotic DSBs is conserved in the mouse. Moreover, we provide the first direct evidence of the interaction between REC114 and MEI4 and identified a potential interaction domain in REC114 that includes previously identified conserved motifs.

### Properties of REC114

Our study revealed that REC114 N-terminus is a PH domain that is composed of two sets of perpendicular anti-parallel β sheets followed by an α helix. This domain is present in a large family of proteins with diverse biological functions and is mostly involved in targeting proteins to a specific site and/or in protein interactions. A subset of these proteins interacts with phosphoinositide phosphates (Lietzke et al. 2000; Lemmon 2003). Several conserved positively charged residues in the two β sheets β1 and β2 important for the interaction are not present in REC114. However, subsequent studies revealed interactions between PH domain and a variety of different partners, in some cases by binding to phosphotyrosine-containing proteins or to polyproline (Scheffzek and Welti 2012). Therefore, the REC114 PH domain could be a platform for several interactions, some of which could involve phosphorylated serine or threonine residues because it has been shown that ATR/ATM signaling through phosphorylation of downstream proteins regulate meiotic DSB activity (Joyce et al. 2011; Lange et al. 2011; Zhang et al. 2011; Carballo et al. 2013; Cooper et al. 2014).

In terms of conservation of the REC114 primary sequence, most of the previously described conserved motifs (SSM1 to 6) are within this PH domain and are readily identified in many eukaryotes. At REC114 C-terminus, the SSM7 motif overlaps with a predicted α helical structure and is less well conserved. Moreover, its presence remains to be established in several species (Tesse et al. 2017). In this study, we demonstrated that this C-terminal domain directly interacts with MEI4, suggesting that this SSM7 region is evolutionarily conserved. The N-terminal domain of MEI4 that interacts with REC114 has a predicted α helical structure and includes two conserved motifs (Kumar et al. 2010).

### Interaction of REC114 with the chromosome axis

A previous study showed that IHO1 is required for MEI4 and REC114 focus formation on axis and that it directly interacts with REC114 by two-hybrid assay (Stanzione et al. 2016). As IHO1 interacts with HORMAD1, IHO1 could act as a platform to recruit REC114 and MEI4. Such a mechanism would be similar the one identified in *S. cerevisiae* for the recruitment of Rec114 and Mei4 by Mer2 (Henderson et al. 2006; Panizza et al. 2011). In agreement with this hypothesis, IHO1 association with the chromosome axis is not altered in the absence of MEI4 or REC114, similarly to what observed in *S. cerevisiae* (Panizza et al. 2011). Therefore, IHO1 could recruit REC114 by direct interaction, and this should allow MEI4 recruitment. Alternatively, we suggest a mechanism where REC114/MEI4 would be recruited as a complex to the axis as we observed a mutual dependency between these two proteins for their axis localization: the formation of REC114 axis associated foci is strongly reduced in the absence of MEI4 and reciprocally. The residual REC114 foci detected in the absence of MEI4 do not appear to be able to promote DSB formation as DSB repair foci are abolished in *Mei4^-/-^* mice similarly to *Spo11^-/-^* mice and thus suggesting an active role for the REC114/MEI4 complex. MEI4 may also be able to interact (directly or indirectly) with IHO1 or with axis proteins independently from REC114 at least in a *Rec114^-/-^* genetic background as weak MEI4 axis associated foci were observed in *Rec114^-/-^* spermatocytes and oocytes and because IHO1 protein was detected upon immuno-precipitation of MEI4 in *Rec114^-/-^* spermatocyte extracts. The details of these interactions and their dynamics during early meiotic prophase remain to be analyzed in details.

Overall, the IHO1/MEI4/REC114 complex is expected to be the main component for the control of SPO11/TOPOVIBL catalytic activity. It may be essential for turning on and off the catalytic activity. In *S. cerevisiae*, it has been proposed that the local control of meiotic DSB formation is constrained by the chromatin loop organization and involves Tel1 (ATM) (Garcia et al. 2015) and possibly also the Mer2/Mei4/Rec114 (IHO1/MEI4/REC114) complex. Indeed, *S. cerevisiae* Rec114 shows Tel1/Mec1-dependent phosphorylation associated with downregulation of DSB activity (Carballo et al. 2013). The IHO1/MEI4/REC114 complex could be a limiting factor for DSB formation. In agreement, we noted that the number of cytologically detectable foci is of the same order (about 200) as the number of DSB events measured by detection of DSB repair proteins. The shutting off of DSB formation that correlates with synapsis between homologs (Thacker et al. 2014) could be the direct consequence of the removal of the Hop1 (or HORMAD1 in mice) axis protein, resulting in the displacement of the Mer2/Mei4/Rec114 (IHO1/MEI4/REC114 in mice) complex from the axis. Additional studies on the protein-protein interactions and post-translational modifications will help to understand these important steps for the regulation of meiotic DSB formation.

## Acknowledgments

We thank Yukiko Imai for many advices on immunoprecipitation assays and Frédéric Baudat for help and advices on histology. We thank all laboratory members for insight and discussions. We thank Amélie Sarazin for image analysis. We thank Attila Toth for comments on the manuscript, and anti-IHO1 and anti-MEI4 antibodies. We thank Morgane Auboiron for help in immuno-cytochemistry. BdM thanks Akira Shinohara and Osaka University for providing support during the preparation of the manuscript. We thank the following BioCampus Montpellier facilities: the Réseau des Animaleries de Montpellier (RAM) for animal care, the Réseau d’Histologie Expérimentale de Montpellier (RHEM) for histology, the Montpellier Resources Imagerie (MRI) for microscopy, and the TAAM/CNRS facility. ABJ was supported by the Labex GRAL (ANR-10-LABX-49-01). This work used the platforms of the Grenoble Instruct-ERIC Center (ISBG: UMS 3518 CNRS-CEA-UGA-EMBL) with support from FRISBI (ANR-10-INSB-05-02) and GRAL (ANR-10-LABX-49-01) within the Grenoble Partnership for Structural Biology (PSB). We thank Caroline Mas and Marc Jamin, for assistance with MALLS and Luca Signor for mass spectrometry analysis. We thank the staff of the ESRF-EMBL Joint Structural Biology Group, particularly Matthew Bowler, for access to and help with the ESRF beamlines. We thank the EMBL high-throughput crystallization facility (HTX). BdM was funded by grants from the Centre National pour la Recherche Scientifique (CNRS) and the European Research Council (ERC) Executive Agency under the European Community’s Seventh Framework Programme (FP7/2007-2013 Grant Agreement no. [322788]). C. O. was funded in part by a post-doctoral fellowship from LabexEpigenMed, program « Investissements d’avenir », ANR-10-LABX-12-01. B.d.M. was recipient of the Prize Coups d’Élan for French Research from the Fondation Bettencourt-Schueller.

## Authors’ contribution

RK initiated the project, analyzed mice and performed cytological analysis

CO analyzed mice and protein interactions

CB analyzed mice, performed cytological analysis and quantified the data

YT prepared samples and antibodies

A-B.J-M performed in vitro assays

JK designed and performed in vitro assays and structural analysis.

BM supervised the project, analyzed the data, prepared figures and wrote the manuscript

## Supplemental figure legends

**Figure S1**

(A) Map of the genomic region including the wild type *Rec114*, *Rec114^-^* and *Rec114^del^* alleles. The six *Rec114* exons are labelled E1 to E6. The conserved motifs are labelled SSM1 to SSM7 (Kumar et al. 2010). Note that SSM2 position was revised by (Tesse et al. 2017). The *Rec114^del^* allele was obtained by expression of the Flip recombinase (Flp) in mice carrying the *Rec114^-^* allele.

(B) Potential proteins encoded by the different *Rec114* alleles

(C) Testis weight is reduced in *Rec114^-/-^* mice compared with *Rec114^+/+^* and *Rec114^+/-^* mice.

Body and testis weights (mean ± SD) were measured in 8-10-week-old *Rec114^+/+^* (testis weight 90±5.7; n=6)*, Rec114^+/-^* (testis weight 94.6±3.6 mg; n=6) and *Rec114^-/-^* (testis weight 25.8±4.3 mg; n=6) males. P values were calculated with the Mann-Whitney two-tailed test.

**Figure S2**

(A) Immunostaining of DMC1 and SYCP3 in oocytes from E15 *Rec114^+/+^* and *Rec114^-/-^* females. Scale bar, 10μm.

(B) Quantification of DMC1 foci (mean ± SD) in leptotene and zygotene oocytes from E15 *Rec114^+/+^* and *Rec114^-/-^* E15 mice (143.0±48.7 and 47.8±27.1 in *Rec114^+/+^* and *Rec114^-/-^* oocytes, respectively; n=55 and 51 what?). P value was calculated with the Mann-Whitney two-tailed test.

(C) Immunostaining of replication protein A2 (RPA2) and SYCP3 in 15 dpp spermatocytes and E15 oocytes from *Rec114^+/+^* and *Rec114^-/-^* mice. Scale bar, 10μm.

(D) Immunostaining of RAD51 and SYCP3 in spermatocytes from 15 dpp *Rec114^+/+^* and *Rec114^-/-^* mice. Scale bar, 10μm.

(E) Immunostaining of SYCP1 and SYCP3 in spermatocytes from 15 dpp *Rec114^+/+^* and *Rec114^-/-^* mice. Scale bar, 10μm.

**Figure S3**

(A) Testis weight is reduced in *Rec114^del/del^* mice compared with *Rec114^+/del^* mice.

Body and testis weights (mean ± SD) were measured in 8-10-week-old *Rec114^+/del^* (testis weight 102.7±13.6 mg; n testis=8) and *Rec114^del/del^* mice (testis weight 31.3±3.9 mg; n testis=8). P values were calculated with the two tailed Mann-Whitney test.

(B) Periodic Acid Schiff staining of testis sections from 9-week-old *Rec114^+/del^* and *Rec114^del/del^* mice. S: Sertoli cell; Sp: Spermatogonia; PL: Pre-leptotene; L: Leptotene; P: Pachytene; Pr-S: Primary spermatocyte; rS: round Spermatid; eS: elongated Spermatid.

(C) Hematoxylin-eosin staining of ovary sections from 9-week-old, *Rec114^+/del^* and *Rec114^del/del^* mice. PF: primary follicle; AF: antral follicle; CL: corpus luteus.

(D) Quantification of primordial, primary, secondary and antral follicles in ovaries from 9-week-old *Rec114^+/del^* and *Rec114^del/del^* mice. P values were calculated with the Mann-Whitney two-tailed test.

(E) Immunostaining of SYCP3, REC114 and γH2AX in leptotene, of SYCP3, RPA and γH2AX in zygotene, and of SYCP3, SYCP1 and γH2AX in zygotene-like or pachytene spermatocytes from 9-week-old *Rec114^del/del^* and *Rec114^+/del^* mice. Scale bar, 10μm.

**Figure S4**

(A) Immunostaining of IHO1 and SYCP3 in early prophase spermatocytes from 13 dpp *Mei4^+/+^* and 12dpp *Mei4^-/-^* males. Scale bar, 10μm.

(B) Immunostaining of MEI4 and SYCP3 in early prophase oocytes from E15 *Rec114^+/+^*, *Rec114^+/-^* and *Rec114^-/-^* females. Scale bar, 10μm.

(C) Quantification of MEI4 foci (mean ± SD) in leptotene oocytes from E15 *Rec114^+/+^* and *Rec114^-/-^* females. The numbers of total foci and foci on chromosome axis (colocalized with SYCP3) were 321±93 and 305±83 in in *Rec114^+/+^* and 88±50 and 82±47 in *Rec114^-/-^* oocytes; n=48 and 47). P values were calculated with the Mann-Whitney two-tailed test.

(D) Quantification of intensity (arbitrary units) of axis-associated MEI4 foci (mean ± SD) in leptotene spermatocytes and oocytes from *Rec114^+/+^* and *Rec114^-/-^* mice: 0.0164±0.0068 and 0.0094±0.0015 in *Rec114^+/+^* and *Rec114^-/-^* spermatocytes (n=12001 and 3775, respectively) and 0.0125±0.0054 and 0.0066±0.0009 in *Rec114^+/+^* and *Rec114^-/-^* oocytes (n=14661 and 3853 respectively). P values were calculated with the Mann-Whitney two-tailed test.

**Figure S5. Comparison of the REC114 PH domain with the N-terminal domain of CARM1.**

(A) Structure of mouse REC114 (15-150).

(B) Structure of CARM1 (2OQB) superimposed to REC114 with r.m.s. deviation of 1.69 Å for 87 Cα atoms.

**Figure S6. Conservation of the REC114 PH domain surface**

(A) Ribbon diagram of the REC114 PH domain.

(B) Surface representation of the PH domain in the same orientation as in A. The conservation of surface residues is represented from grey to green, according to the color scale shown below and based on the sequence alignment shown in Fig. 6D. The position of the most conserved and exposed residues is shown.

(C) Ribbon diagram of the PH domain, but rotated 180° around the vertical axis, compared with A.

(D) Surface conservation of the PH domain corresponding to the view shown in C.

**Figure S7. Dimerization of Rec114**

(A) The REC114 PH domain forms dimers with a symmetry-related molecule. The dimer interface includes the α1 helix and β6 strand. The key interacting residues are indicated.

(B) Molecular mass determination of the mouse REC114 PH domain by multi-angle laser light scattering (MALLS). 50 μL of purified protein at a concentration of 10, 5 and 2 mg/ml was injected in a Superdex 75 gel filtration column. Online MALLS and refractive index data were recorded on a DAWN-EOS detector (Wyatt Technology Corp.) using a laser emitting at 690 nm and an Optilab T-rEX detector (Wyatt Technology Corp.) respectively, with a refractive-index increment dn/dc of 0.185 mL.g^-1^. Data were analyzed using the ASTRA 6 software (Wyatt Technology Corp.). The molecular mass could only be determined for samples injected at 10 and 5 mg/ml (MW in light and dark blue, respectively). The results indicate that REC114 exists in solution as a concentration-dependent mixture of dimers and monomers with fast exchange rate.

## Material and Methods

### Mouse strains

The non-conditional *Rec114* mutant allele is referenced as 2410076I21Rik^tm1(KOMP)Wtsi^ and named *Rec114^-/-^* in this study. The *Rec114^del^* allele, in which the inserted lacZ-Neo cassette was deleted, was obtained by expression of Flp in mice carrying *Rec114^-/-^*. These mice are in the C57BL/6 background. The *Mei4^-/-^* and *Spo11^-/-^* strains were previously described (Baudat et al. 2000; Kumar et al. 2010). All animal experiments were carried out according to the CNRS guidelines and approved by the ethics committee on live animals (project CE-LR-0812 and 1295).

### Antibodies

Chicken anti-REC114 antibodies were generated against mouse REC114 and affinity-purified. The anti-SYCP3, anti-MEI4 and anti-IHO1 antibodies were previously described (Baudat and de Massy 2007; Kumar et al. 2010; Stanzione et al. 2016). Other antibodies used in this study were against γH2AX (Millipore 05-636), DMC1 (Santa Cruz, SC-22768), SYCP1 (Abcam ab15090), RPA32 (gift from R. Knippers) and RAD51 (gift from W. Baarends). For immunofluorescence experiments, the following secondary antibodies were used: Cy™3 AffiniPure Donkey Anti-Guinea Pig IgG (H+L) (Jackson ImmunoResearch), Donkey anti-Rabbit IgG (H+L) Alexa Fluor 488 (Thermo Fisher Scientific) and Donkey anti-Mouse IgG (H+L) Alexa Fluor 647 (Thermo Fisher Scientific).

### Histology and cytology

Testes and ovaries were fixed in Bouin’s solution (Sigma) at room temperature overnight or for 5 h, respectively. After dehydration and embedding in paraffin, 3-μm sections were prepared and stained with Periodic acid-Schiff for testis and with hematoxylin and eosin for ovaries. Image processing and analysis were carried out with the NDP.view2 software (Hamamatsu).

Spermatocyte and oocyte chromosome spreads were prepared by the dry-down method (Peters et al. 1997).

### Image analysis

γH2AX was quantified using Cell profiler 2.2.0. The total pixel intensity per nucleus was quantified. The intensity of MEI4 foci was the mean pixel value within a focus. Axis-associated MEI4 foci were determined by co-labelling with SYCP3.

### Protein analysis

Whole testis protein extracts were prepared as described in (Stanzione et al. 2016).

For REC114 immunoprecipitation, 1.5mg of extract was diluted in IP buffer (20mM Tris-HCl; 150mM NaCl; 0.05% NP40; 0.1% Tween-20; 10% glycerol; protease inhibitors) and incubated with 2 μg of affinity-purified chicken anti-REC114 antibody at 4°C overnight. Then, 50 μl of agarose-immobilized anti-chicken IgY Fc (goat) (GGFC-130D, Icllab) was added at 4°C for 1h. Beads were washed five times with washing buffer (20mM Tris-HCl pH 7.5, 0.05% NP-40, 0.1 % Tween-20, 10% glycerol, 150mM NaCl). Immunoprecipitated material was eluted and incubated with 2X Laemmli loading buffer (with 20mM DTT) at 95°C for 5 min.

For MEI4 immunoprecipitation, 3 μg of guinea pig anti-MEI4 antibody (Stanzione et al. 2016) was crosslinked to 1.5 mg of Dynabeads Protein A (Invitrogen) with disuccinimidyl suberate using the Crosslink Magnetic IP/Co-IP kit (Pierce, ThermoFisher Scientific). 3.6 mg of testis protein extract was incubated with the crosslinked antibody at 4°C overnight. Beads were washed five times with washing buffer (20mM Tris-HCl pH 7.5, 0.05% NP-40, 0.1 % Tween-20, 10% glycerol, 150mM NaCl). Immunoprecipitated material was eluted by incubating the beads with the Elution Buffer (pH 2) for 5min, and neutralized with the Neutralization Buffer (pH 8.5) (both buffers provided with the kit). Eluates were incubated with Laemmli loading buffer (1X final) at RT for 10min, and divided in three aliquots, adding 10mM DTT to one of them (for REC114 detection), followed by incubation at 95°C for 5min.

### Western blot analysis

Immunoprecipitates and inputs were separated on 10% Mini-PROTEAN TGX Precast Gels (Bio-Rad) and then transferred onto nitrocellulose membranes with the Trans-Blot Turbo Transfer System (Bio-Rad). The following primary antibodies were used: rabbit anti-MEI4 (1/500; Kumar et al., 2010), chicken anti-REC114 (1/1000), rabbit anti-IHO1 (1/2000; Stanzione et al., 2016), and guinea pig anti-SYCP3 (1/3000; Kumar et al., 2010). The following horseradish peroxidase (HRP)-conjugated secondary antibodies were used: anti-rabbit (1:5000; Cell Signaling), True-Blot anti-rabbit (1/1000, Jackson ImmunoResearch), donkey anti-chicken IgY (1/3000; Jackson ImmunoResearch), and donkey anti-guinea pig (1/10000; Jackson ImmunoResearch).

#### Protein expression, purification and crystallization

Mouse REC114 (15-159) fused to His-tag was expressed in *E. coli* BL21-Gold (DE3) (Agilent) from the pProEXHTb expression vector (Invitrogen). The protein was first purified by affinity chromatography using Ni^2+^ resin. After His-tag cleavage with the TEV protease, the protein was further purified through a second Ni^2+^ column and by size-exclusion chromatography. Pure protein was concentrated to 10 mg.ml^−1^ in buffer (20mM Tris, pH 7.0, 200mM NaCl and 5mM mercaptoethanol). The best-diffracting crystals grew within 1 week at 20°C in a solution containing 0.25M ammonium sulfate, 0.1M MES (pH 6.5) and 28% PEG 5000 MME. For data collection at 100 K, crystals were snap-frozen in liquid nitrogen with a solution containing mother liquor and 25% (v/v) glycerol. SeMet-substituted REC114 was produced in *E. coli* BL21-Gold(DE3) and a defined medium containing 50 mg.l^−1^ of SeMet. SeMet REC114 was purified and crystallized as for the native protein.

#### Data collection and structure determination

Crystals of REC114 (15-159) belong to the space group *P*6_1_22 with the unit cell dimensions *a*, *b* = 107.5 Å and *c* = 82.8 Å. The asymmetric unit contains one molecule and has a solvent content of 71%. A complete native dataset was collected to a resolution of 2.5 Å on the beamline ID30A-1/MASSIF-1 at the ESRF (Grenoble, France). The SeMet REC114 crystallized in the same conditions in the space group *P*4_2_2_1_2 and contained two molecules per asymmetric unit. A complete SeMet dataset was collected to a resolution of 2.7 Å at the peak wavelength of the Se K-edge on the ID23-1 beamline at the ESRF. Data were processed using XDS (Kabsch 2010). The structure was solved using SeMet SAD data. Selenium sites were identified, refined and used for phasing in AUTOSHARP (Bricogne et al. 2003). The model was partially built with BUCCANEER (Cowtan 2006), completed manually in COOT (Emsley et al. 2010) and refined with REFMAC5 (Murshudov et al. 1997). The model was used for molecular replacement to determine the structure using the native dataset and PHASER (McCoy et al. 2007). The native structure was finalized in COOT and refined with REFMAC5 to a final *R*-factor of 25% and *R*_free_ of 30% (Table 1) with all residues in allowed regions (96% in favored regions) of the Ramachandran plot, as analyzed using MOLPROBITY (Chen et al. 2010).

#### Strep-tag pull-down assays

MEI4 (1-127) was cloned in the pProEXHTb expression vector to produce a His-tag fusion protein. REC114 and its deletion mutants were cloned in the pRSFDuet-1 vector as Strep-tag fusion proteins. REC114 variants alone or co-expressed with MEI4 were purified using a Strep-Tactin XT resin (IBA). The resin was extensively washed with a buffer containing 20mM Tris, pH 7.0, 200mM NaCl and 5mM mercaptoethanol, and bound proteins were eluted with the same buffer containing 50mM biotin and analyzed by 15% SDS-PAGE. The minimal REC114-MEI4 complex was then purified using the Strep-Tactin XT resin. The His-tag of MEI4 was removed with the TEV protease and a passage through a Ni^2+^ column. The complex was then purified by size-exclusion chromatography.

